# A behavioral paradigm for measuring perceptual distances in mice

**DOI:** 10.1101/2021.04.29.441965

**Authors:** Hirofumi Nakayama, Dmitry Rinberg

## Abstract

Perceptual similarities between a specific stimulus and other stimuli of the same modality provide valuable information about the structure and geometry of sensory spaces. While typically a subject of behavioral experiments in humans, perceptual similarities, or distances, are rarely measured in other species. However, understanding the neural computations responsible for sensory representations requires the monitoring and sometime manipulating of neural activity, which is more readily achieved in other experimental models. Here we develop a behavioral paradigm that allows for the quantifying of perceptual similarity between sensory stimuli using mouse olfaction as a model system.

---

Perceptually similar objects are represented by similar patterns of neuronal activity in the brain [1]. Thus, for multiple sensory objects, both their relative perceptual distances and the structure of their perceptual space--its dimensionality and topology--should be reflected in the corresponding space of their neural representations. For example, the perceptual proximities between different spectral colors reveals the circular organization of hue in two-dimensional space [2, 3], and the perceptual similarity between different sounds led to the discovery of the helical organization of musical tone [4]. Such information about perceptual space is indispensable for the understanding of neural coding mechanisms underlying sensory perception.

However, it is usually difficult to perform both perceptual and neural measurements in the same model organism. Most attempts to reveal the perceptual relationship between stimuli were performed in psychophysical studies with either human or monkey subjects (but see [5, 6]). Recent technological progress makes the mouse an attractive model for recording [7, 8] and manipulating [9–13] neural activity at different temporal and spatial scales. However, a lack of behavioral methods for measuring perceptual distances prevents the linking of neural activity with perceptual space in the same species.

Here we develop a robust and high-throughput method for estimating perceptual distances between multiple pairs of sensory stimuli in mice using olfaction as a model system. The choice of olfaction is dictated by two main factors. First, olfaction is a highly relevant sensory modality for rodents, which makes it easier to train animals on demanding behavioral paradigms. Second, olfaction is one of the least explored primary sensory systems, and our understanding of the structure of olfactory perceptual space is still limited [14–16]. Individual odorants (chemical molecules) are characterized by thousands of physicochemical features, and the majority of stimuli experienced by animals in the wild are complex odor mixtures. How multidimensional sensory space is represented in the brain remains unknown.

In human psychophysics, odor perceptual space has been characterized using multiple approaches including semantic descriptors [17–19], analog ratings of perceptual qualities [20, 21], and analog ratings of perceptual similarity between odors pairs [22]. Among them, only similarity ratings between odors can be obtained in non-human model animals including rodents. Given a set of pairwise similarity (or distance) of perception across odors, dimensionality reduction algorithms lead to the interpretable low dimensional structure that represent perceptual quality of odors. In rodent experiments, the cross-habituation paradigm has been used to assess if pairs of odors are perceptually similar or not [23]. However, the paradigm is low-throughput and requires many animals for each of odor pair. Due to this limitation, it is challenging to collect sufficient pairwise similarity measurements to recover the structure of perceptual space.

To overcome this limitation, we designed a behavioral paradigm that allows us to perform high-throughput measurement of perceptual distances across many odor pairs. We used a delayed match-to-sample (DMTS) behavioral paradigm, where mice are trained to respond differently depending on whether two sequentially presented odors are the same or different. The probability that a trained mouse responds as if two odors within a trial are the same (or different) is taken to reflect odor pairwise perceptual similarities (or distances).

The DMTS paradigm was originally developed for pigeons and monkeys [24, 25], and has been used to study short-term memory in primates and, more recently, visual perception [26]. The olfactory DMTS task was initially established for freely moving rodents [27] and recently used with head-fixed rodents to study short-term memory [28–30]. In these experiments, only a small number of stimuli (usually two odors) were used. Introducing every new odor pair would require new training and testing many odor pairs would be impractically time consuming [30].

Differently from previous studies, we trained mice to perform the task against many odors, thus preventing mice from forming behavioral strategies that are only applied to specific odors. This allows us to use the task for high-throughput measurement of pairwise odor perceptual distances--similar to that undertaken in human studies. We collected behavioral data for more than a hundred of odor pairs, and demonstrated the relationship between a newly proposed behavioral measurement and perceptual odor similarities.

The behavior paradigm was designed to make animals compare two odors that are presented within the same trial. Mice received two sequential odor presentations (1 s duration each, and 5 s inter-stimulus interval) followed by a 1 s response window (**Fig. 1a**). If two odor stimuli were the same (a match trial), mice should respond by licking a center water spout during the response time window (go response). If not (a non-match trial), the correct response was to suppress licking (nogo response) (**Fig. 1b**). In non-match trials, mice were rewarded for no-go responses with a drop of water from a side water spout. Although two water spouts are used, licking or not-licking the side water spout does not affect the reward contingency. Thus, we still consider this paradigm to be based on go/no-go decisions.

**Figure 1.**
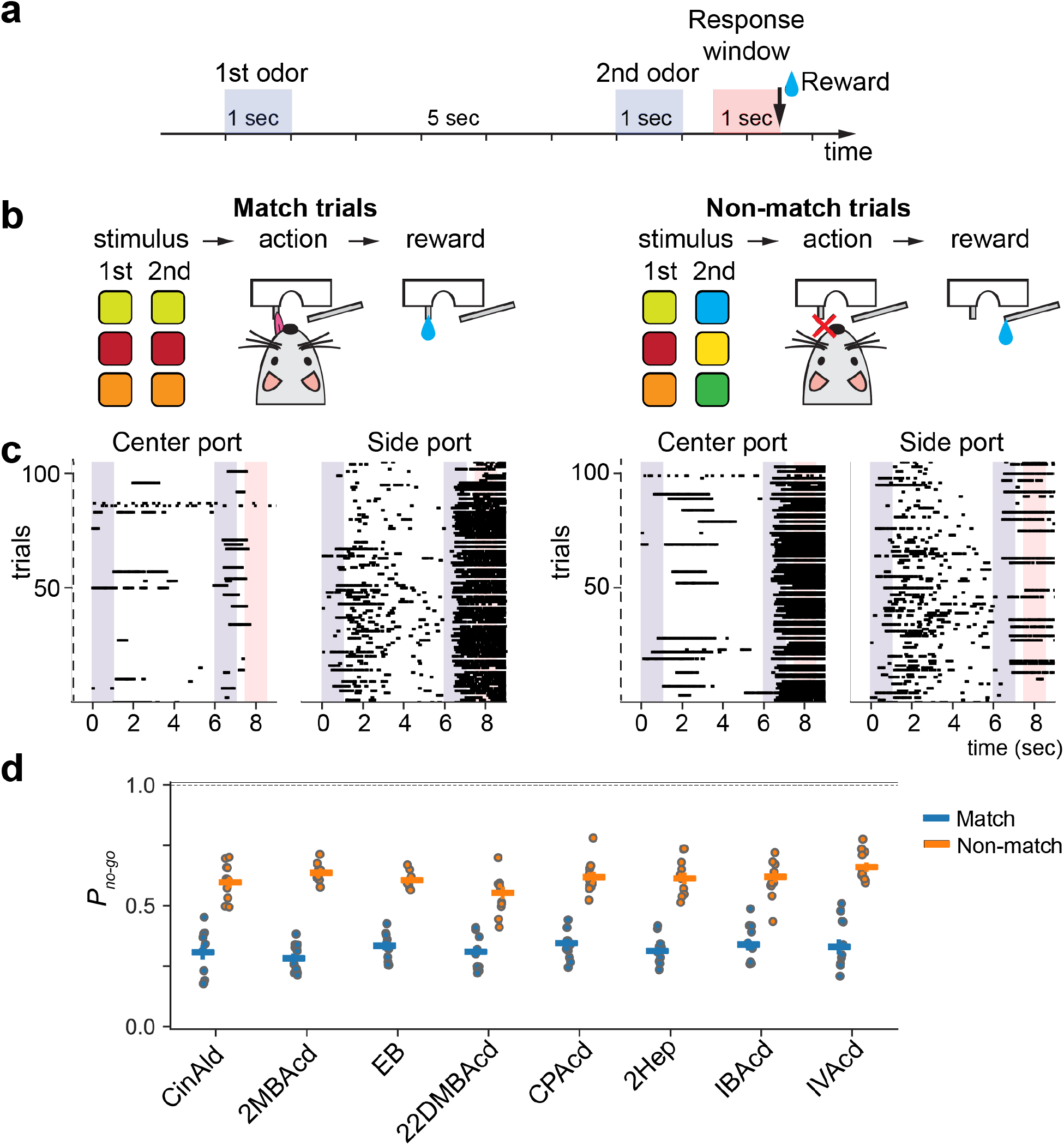
Characterization of task performance. **a**, The sequence of events in a delayed match-to-sample task trial. **b**, Schematics of two trial types. *Left panel:* match trials - the correct behavior response is licking a center port (go), followed by a water reward provided from the same port. *Right panel:* non-match trials - the correct behavior response is not licking the center port (no-go), followed by a reward provided from a side lick port. Licking to the side port during the response window did not affect outcomes. **c**, Lick patterns in example trials. Each row represents a single trial. Licks are shown as black ticks. **d**, Probability of a nogo choice by odor identity. Trials were assigned to a specific odor identity if that odor was presented either on first or second presentations. Circles correspond to individual mice, bars are averages across mice.

Odors in each trial were pseudorandomly chosen from a panel of monomolecular odors and their binary mixtures (**Table 1**). To discourage mice from comparing two odors based on their intensity, we randomized odor concentrations at each trial for the same odor identity presenting either high or low concentrations with 5-fold differences.

**Table 1.** Odors used in the experiments.

| Set | Odor name | Abbreviation | Liquid dilution | Headspace concentration (M) | Air dilution range |
| --- | --- | --- | --- | --- | --- |
| 1 | Cinnamaldehyde | CinAld | 0.0004 | $8.9 \times 10^{-10}$ | 0.01 – 0.05 |
| | Ethyl butyrate | EB | 0.0008 | $9.1 \times 10^{-7}$ | |
| | 2-Methylbutyric acid | 2MBAc | 0.0004 | $8.9 \times 10^{-9}$ | |
| | 2,2-Dimethylbutyric acid | 22DMBAc | 0.0004 | $2.3 \times 10^{-9}$ | |
| | Cyclopentanecarboxylic acid | CPAc | 0.0012 | $1.1 \times 10^{-8}$ | |
| | 2-Heptanone | 2Hep | 0.0002 | $4.2 \times 10^{-8}$ | |
| | Isobutyric acid | IBAc | 0.0004 | $7.6 \times 10^{-8}$ | |
| | Isovaleric acid | IVAc | 0.0004 | $1.6 \times 10^{-8}$ | |
| 2 | 3-Heptanone | 3Hep | 0.001 | $1.8 \times 10^{-7}$ | 0.02 – 0.1 |
| | Methylvalerate | MVT | 0.0002 | $1.6 \times 10^{-7}$ | |
| | Ethyl butyrate | EB | 0.0018 | $2.0 \times 10^{-6}$ | |
| | Propionic acid | PpAc | 0.0063 | $2.0 \times 10^{-7}$ | |
| | Butyric acid | ButAc | 0.001 | $4.5 \times 10^{-8}$ | |
| | (+)-α-Pinene | Pinene | 0.001 | $2.9 \times 10^{-7}$ | |
| | Benzaldehyde | BzAld | 0.0004 | $4.9 \times 10^{-8}$ | |
| | 5-Methyl-2-Hexanone | 5M2H | 0.0002 | $6.9 \times 10^{-8}$ | |
| 3 | 3-Heptanone | 3Hep | 0.001 | $1.8 \times 10^{-7}$ | 0.01 – 0.05 |
| | Ethyl butyrate | EB | 0.0018 | $2.0 \times 10^{-6}$ | |
| | Valeric acid | ValAc | 0.0002 | $3.5 \times 10^{-9}$ | |
| | 3-Methylvaleric acid | 3MVAc | 0.0002 | $2.3 \times 10^{-9}$ | |
| | 3,3-Dimethylbutyric acid | 33DMBAc | 0.0002 | $4.6 \times 10^{-9}$ | |
| | (+)-α-Pinene | Pinene | 0.001 | $2.9 \times 10^{-7}$ | |
| | Benzaldehyde | BzAld | 0.0004 | $4.9 \times 10^{-8}$ | |
| | Isovaleric acid | IVAc | 0.0002 | $7.8 \times 10^{-9}$ | |
| | 4-Methylvaleric acid | 4MVAc | 0.0002 | $2.1 \times 10^{-9}$ | |
| | Hexanoic acid | HexAc | 0.0002 | $2.8 \times 10^{-9}$ | |

Trained mice exhibited differential lick responses between match and non-match trials (**Fig. 1c**) with the probability of a no-go response being 0.31 ± 0.02 and 0.65 ± 0.04 respectively (mean ± std, across all odor identities and all mice) (**Fig. 1d**).

We collected data from 10 mice over 308 sessions (75195 total trials after excluding the first 5 trials of each session) for a panel of 8 monomolecular odors and 16 binary mixtures (**Fig. 2a**). The probability of an error in non-match trials varied across different odor pairs, and we assumed it may be a measure of odor perceptual similarity. Thus, we propose a distance metric as the probability of correct responses in non-match trials *(P_no-go_)* normalized by the probability of error responses on match trials for identical odors in a way similar to previous work [31]:

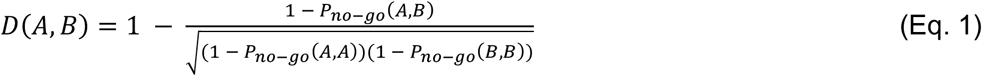

**Figure2.**
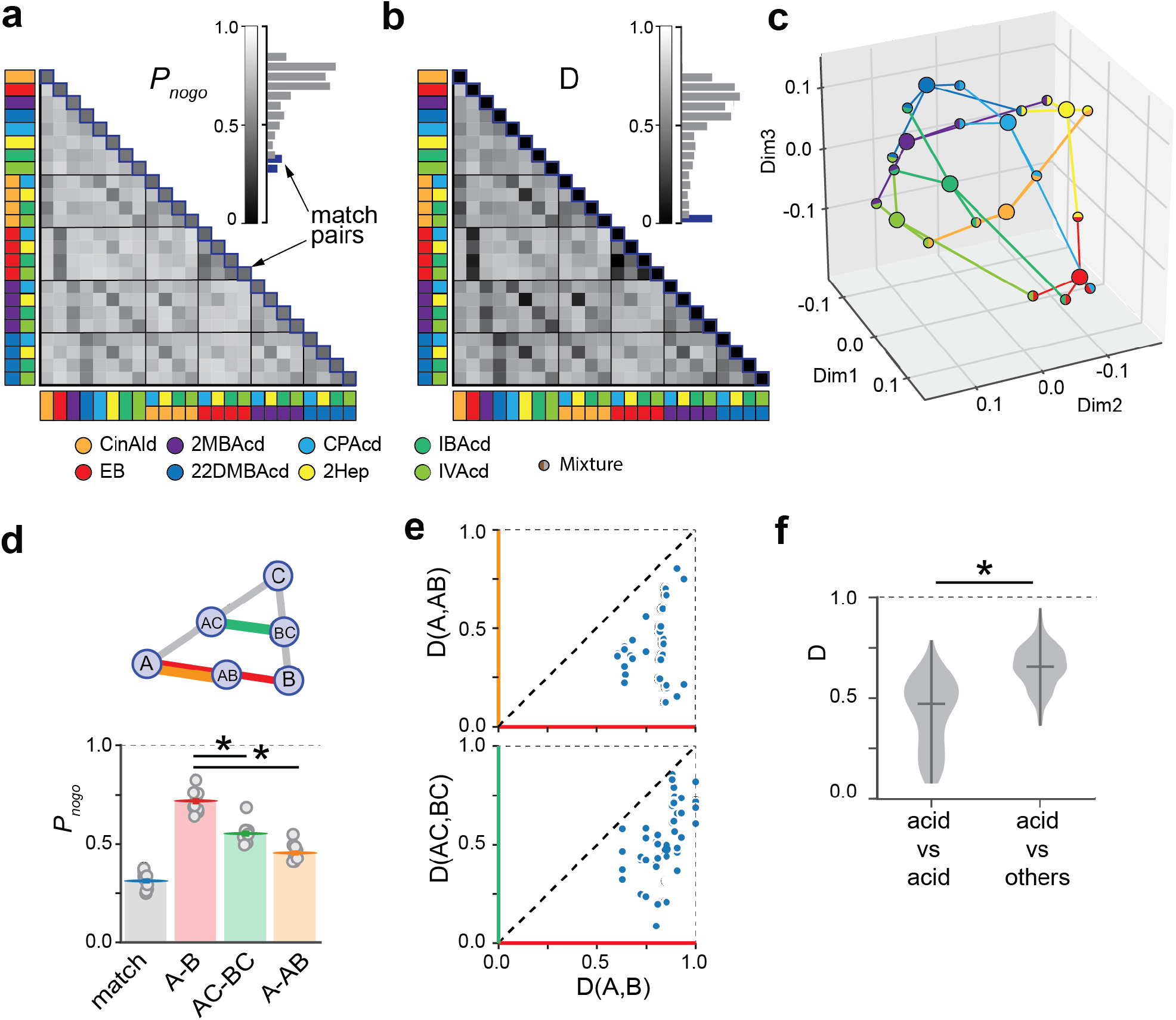
Validation of the behavioral readout as a measure of a perceptual distance. **a**, Matrix and distribution (insert) of probabilities of a no-go choice for each of odor pair. **b**, Perceptual distance matrix calculated from the matrix in **a. c**, MDS embedding of odors calculated from the distance matrix in **b. d**, *Top:*A schematic illustrating perceptual distances between binary mixtures and their component odors. *Bottom:*Average probabilities of a no-go choice for match trials and three types of non-match trials corresponding to different odor and binary mixture trials. **e**, Individual odor pair scatter plots corresponding to the relationship shown in **d**, *top: D(A, AB)* vs *D(A, B), bottom: D(AC, BC)* vs *D(A, B).***f**, Distribution of the distances between acids (mean: *D_median_*= 0.47) and between acids and non-acids (mean: *D_median_*= 0.66).

The distance is equal to zero for identical odors, and increases as odors became less similar. The distribution of the distances for all measured odor pairs is presented in **Fig. 2b**. For data visualization purposes, we applied multidimensional scaling (MDS) [32–34] to the distance matrix and presented an odor-odor distance graph in the three-dimensional space (**Fig. 2c**).

To test the assumption that our proposed distance metric reflects a perceptual distance between odors, we performed a series of experiments with binary mixtures. For humans, the most common outcome of perceiving a binary mixture is an average of the qualitative odor descriptors for the component molecules [35]. Thus, we first hypothesized that the distance between two odors should be greater than between one of them and their binary mixture: *D(A,AB) < D(A,B).* Second, we assumed that the distance between two binary mixtures sharing a common component should be lesser than the distance between odors not sharing a common component: *D(AC,BC)*<*D(A,B)* (**Fig. 2d**). We find that both of these assumptions are satisfied, on average (**Fig. 2d**) and for each odor pair (**Fig. 2e**). *(D(A,B) > D(A,AB): p*= 0.0039, *D(AC,BC) < D(A,B): p*= 0.0039, n = 10 mice, Wilcoxon signed-rank test after Bonferroni correction of *p*-values.)

From human studies we also know that chemically similarity, on average, correlates with perceptual similarity [22]. To test this in our data, we divided odors into two groups: acids and non-acids. The average perceptual distance between acids was significantly smaller than for between acids and non-acids (median distance within acids (163 pairs): 0.47, median distance between acids and non-acids (220 pairs): 0.66 (*p* < 10^−33^, Mann-Whitney U-test) (**Fig. 2f**).

Our analyses of binary mixtures and odors from different chemical groups provide encouraging evidence that our metric can be used to measure distances in perceptual space. We further performed a series of experiments and analyses to confirm that the behavioral responses primarily depend on odor identity rather than other aspects of the task including odor concentration, shortterm memory, long-term drift in performance, and odor presentation sequence.

To investigate the influence of different variables on choice behavior, we fit logistic regression models, which was designed to predict choice based on the distance metric, *D(A,B),* and other behavioral variables such as 1) concentrations of the 1^st^ and 2^nd^ odor in the trail: *X_conc1_, X_conc2_= 0 or 1* for the low or high concentrations; 2) phase of data collection, *X_trials_=* 0 *or* 1 for the 1^st^ vs 2^nd^ half of total set of trials; and 3) a sequence of odor presentation, *X_seq_*= −1 *or 1* for A->B and B->A odor presentations in non-match trials and *X_seq_=* 0 for match trials. To compare the regression coefficients, we normalized the variances and subtracted the means of individual independent variables. The absolute value of regression coefficients in the model indicated that the distance metric had larger influence on choice behavior than other task variables: *D(A,B)* vs *X_conc1_, p* < 0.001; *D(A,B)* vs *X_conc2_*, *p* < 0.001; *D(A,B)* vs *X_trails_, p* < 0.001; *D(A,B)* vs *X_seq_*, *p* <0.001 (Tukey’s honestly significant difference test) (**Fig. 3a**). As expected from the regression coefficient, logit(*P_nogo_*) (the left-hand side of (Eq.2)) increased as the distance metric *D(A,B)* increases (**Extended fig. 2a**). In a separate experiment we vary the delay between odor presentations *(X_delay_*= 0 or 1, for 3 s and 5 s delays) and repeated regression analysis for a new data set adding the additional independent variable *X_delay_*. Again, we found that the effect of distance metric is significantly large than that of the delay: *D(A,B)* vs *X_delay_*, *p* < 0.001 (**Fig. 3b, Extended fig. 2b**).

**Figure 3.**
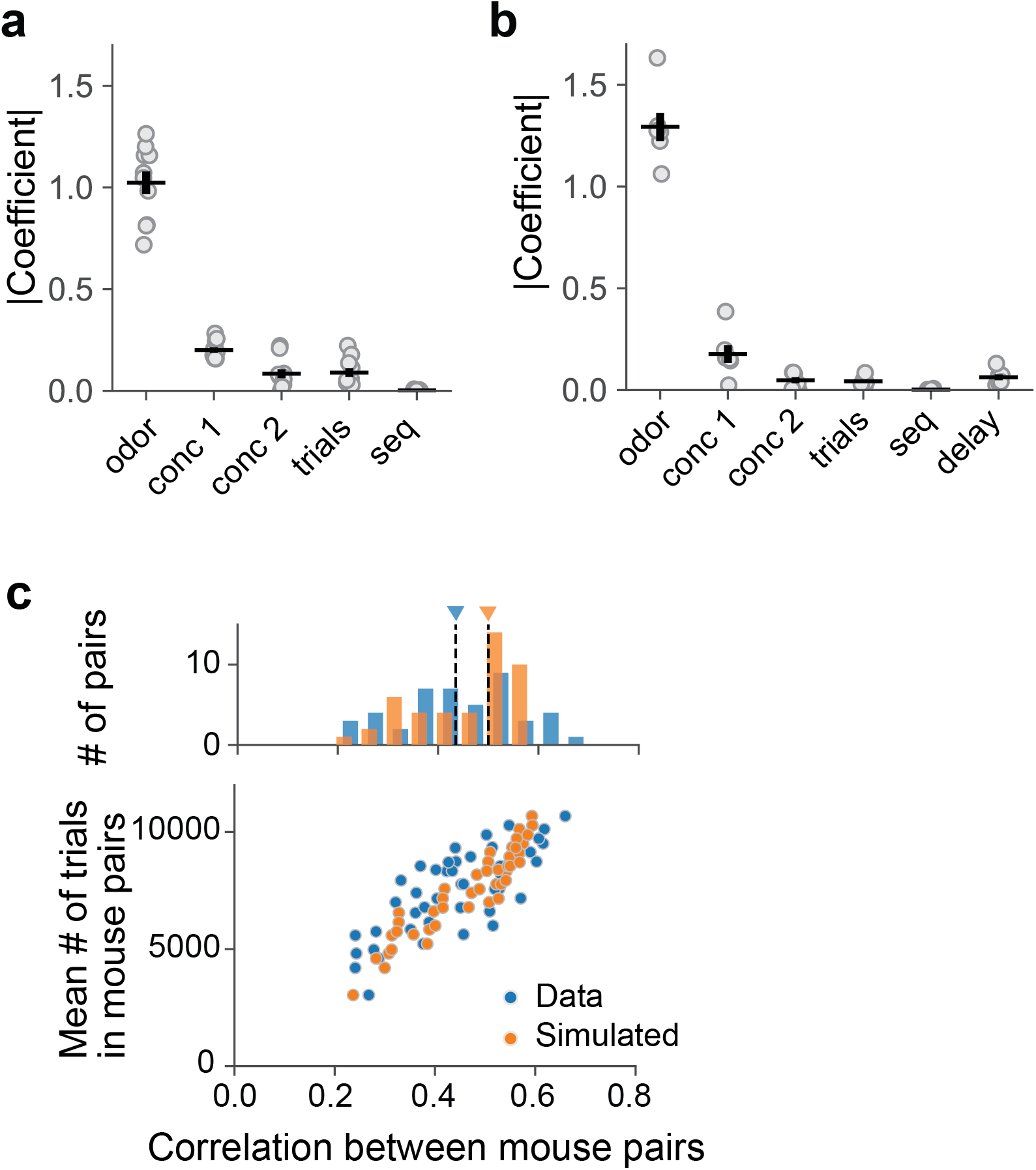
Odor pair dependence and across animal conservation of behavioral readouts. **a**, Absolute value of regression coefficients in the regression model (**a**) and relationship between distance metric and left-hand side of **Eq.2**(**b**), Same as in **a** and **b** for regression model including variable delay **Eq.3**. **c**, Comparison of perceptual distances across animals. *Top*: Distribution of correlation coefficients of perceptual distances between mouse pairs. *Bottom:* The relationship between the number of trials and the pairwise correlation of perceptual distances between mouse pairs.

All experiments indicate that the proposed metric reflects perceptual distances, similar to what is undertaken in human experiments. The critical question remains if the measurement is conserved across mice. Potentially, each mouse may exploit a single perceptual feature or subset of features to make judgements in the DMTS paradigm, and mice may not share the same criteria. We found that correlations across mice are relatively high: a median Pearson correlation coefficient equal to 0.44, higher than values previously reported in humans [21]. Moreover, the correlation coefficient obtained from data was not significantly different from that calculated from simulated data assuming a common underlying perceptual distance matrix for all mice (**Fig.3e, top**, median correlation in data (45 pairs): 0.44, median correlation in simulated data (45 pairs) 0.51, p = 0.077, Wilcoxon signed-rank test). Finally, this coefficient also increases with the number of trials (**Fig. 3e, bottom**), indicating that the correlation of perceptual distances across mice would increase with a greater number of trials.

These experiments indicate that the behavioral readouts are consistent with what is expected from perceptual distances. Notably, we found that perceptual distances largely agree across different animals. This observation, along with the possibility of testing many odors, makes our paradigm a promising tool to investigate olfactory perceptual space. Robust psychophysical measurements of the perceptual distances in mice allows for capitalizing on all advances in mouse genetics and modern methods for recording and manipulating neural activity. In addition, these methods have a lower implementation threshold in head-fixed animals. We expect our behavioral paradigm will contribute to the study of the neural coding of olfaction through direct comparison between perceptual evaluation of odors and corresponding neuronal activity. Beyond research into mouse olfaction, this paradigm allows sensory perception to be studied in generalized settings similarly to human psychophysics.

## Methods

### Animals

Both male and female Thy1-GCaMP6f (GP5.11) mice (Stock No: 024339) (Jackson laboratories) were used in the task. At the start of behavioral training, mice were at least two months old and had 20 g body weight. Mice were housed under a 12-hour inverted light/dark cycle. All procedures were approved under a New York University Langone Health institutional animal care and use committee (IACUC).

### Animal surgeries

Prior to behavioral training, mice were implanted with a 3D-printed head-bar for head fixation in the behavioral apparatus. Mice were anesthetized with isoflurane (2% for induction, 1.5% during surgery) and placed on a heated floor during surgery. Skin overlying the skull was sterilized with betadine and incised to expose the skull. The periosteum was gently scraped away and the surface of the skull was cleaned with hydrogen peroxide. The head-bar was fixed to the skull using C&B Metabond dental cement (Parkell).

### Odor delivery

A two-cassette air-dilution olfactometer was used to prepare and deliver odors with specific concentrations (**Extended Fig. 1**). Each olfactometer cassette consisted of two mass flow controllers (MFCs), (Alicat, MC-100SCCM-D/5M/5IN and MC-1SLPM-D/5M/5IN), four inline teflon four-valve manifolds, (NReserach, 225T082), one on-off clean-air three port bypass valve (NResearch, TI1403270), and eight odor vials. Odors were diluted in water and stored in amber volatile organic analysis vials (Restek, 21797). The total air flow (usually 1000 ml/min) and relative odor concentration were controlled by MFCs. To deliver the odor stimuli, specific odor valves were opened prior to the beginning of each trial and the odorized air flow was diverted to the exhaust line by a final valve (NResearch, SH360T042), while a controlled flow of clean air at the same flow rate was delivered to an odor port. After flow stabilized (~1 sec), the final valve switches between odorized flow and clean air flow, and odor is delivered to the odor port with <100 ms latency. At the end of stimulus presentation, the final valve switches again and delivers clean air to the odor port. The olfactometer enables the dilution of odors between 10- and 100-fold, as well as the delivery of binary mixtures by mixing two regulated flows from different cassettes. The binary mixture ratio was controlled by relative flows through odor vials, while total flow was kept constant.

### Odor stimulus

Eight or ten monomolecular odors and their binary mixtures were used for each experiment **(Table 1)**. All monomolecular odors were diluted in deionized water with following dilutions in v/v and stored in vials during experiments. These odor dilutions were prepared daily and were placed in two different cassettes. Each olfactometer contained four or five odors to enable the generation of binary mixtures by mixing odors from different cassettes. The initial headspace concentration was estimated based on saturated vapor pressure from a given chemical at a specific dilution. Following this, the odor was diluted between 10- and 100-fold using air dilution in the olfactometer. Specific dilutions for each experiment are presented in **Table 1**.

### Behavioral setup

Mice were head-fixed and placed on a wheel allowing free running and experienced minimal stress (**Extended Fig. 2**). A mouse snout was placed into a specially designed odor port (made of PTFE). The odor port was connected to the olfactometer, from which odorized air was delivered, and a vacuum line was present to quickly remove the odorized air during interstimulus intervals. Mice were trained to register their behavioral judgments by licking water spouts, and licking was detected using a capacitive touch sensor (Sparkfun, SEN-1204). Two water spouts were installed, one in the center and the other to the side of the position of a mouse snout in the odor port. Water delivery was controlled by a pinch valve (Valcor, SV74P61T). All behavioral events (stimulus delivery, water delivery, and lick detection) were monitored and controlled by custom programs written in Python interfacing with a custom behavioral control system (Janelia Research Campus) based on an Arduino Mega 2560 microcontroller.

### Behavioral paradigm

The behavioral paradigm aims to train mice to judge whether two odors presented in each trial are the same or not. In each trial, mice were presented with two sequential 1 s odor stimuli, separated by a delay period of 5 s. The odor stimuli were either the same (match trails) or different (non-match trails). Mice were trained with a lick based go-nogo paradigm and instructed to report their behavioral judgment during a 1 s response window, which occurs 0.5 – 1.5 s after the offset of the second odor presentation in each trial (**Fig. 1 a-b**). Odor concentration for each presentation was randomized by adjusting the air flow. Only the center water spout was used to evaluate behavioral response, and licking the side water spout had no effect on trial outcome. Mice were trained to lick (go choice) the center water spout in match trials and to suppress licks (no-go choice) in non-match trials. At the end of a response window, correct responses were followed by reward delivery in the center spout (go choice in match trials) or in the side water spout (no-go choice in non-match trials).

### Behavioral training

The training procedure consisted of multiple steps outlined below:

#### 1. Water restriction and habituation

During the entire period of behavioral training, mice were kept under water deprivation, which began at least one week after surgery. Mice received a total of 1 ml of water per day either as accumulated rewards for behavioral performance, or supplemented to the full amount at the end of the day. During the initial phase, mice were habituated to the handling by experimenters for 10 – 15 min per day. Once mice comfortably drink water drops on the glove of experimenter’s hand, mice began being habituated to the experimental setup. During the habituation to experimental setup, mice were head-fixed on the running wheel and placed in the behavioral box for 10-15 min per day. During this phase, mice were occasionally presented with water through a syringe. Once mice show reliable licking responses to water delivered through a syringe, mice began lick training.

#### 2. Lick training

The purpose of this phase was for mice to learn to lick water spouts to obtain a water reward. Mice were given a 1 - 1.5 ul water drop each time they licked water spouts. Both center and side water spouts were introduced and a water reward was provided alternately from them. In case mice did not lick the water spouts at all, multiple water drops were delivered by experimenters until mice started licking water spouts. Mice proceeded to the next training step if they received 100 water drops within 30 minutes. Otherwise, lick training was repeated in the following days.

#### 3. Behavioral shaping step one - Pavlovian shaping

The purpose of this step is to get mice familiarized with the timing of behavioral events and water reward delivery. At each trial, mice were presented with the same odor twice with 3 s delay, which was followed by a 1 s response window (0.2 – 1.2 s from the second odor offset). At this step, only match trials were presented and two odor presentations in a given trial were always the same. 5 ul of water reward were delivered at the end of response window regardless of behavioral responses. Although licking behavior of mice had no consequence on reward delivery, mice started to show anticipatory licking during the response time window. If mice licked during the response window at >80% of trials across 80 trials, mice proceeded to the next step. In shaping steps one to three, all odors in odor set 2 in Table 1 were used.

#### 4. Behavioral shaping step two – odor lick association

The purpose of this step was to make mice lick during the response window to receive a water reward. The trial structure was same as in step one, except mice had to lick a water spout during the response window (0.5 – 1.5 sec from the second odor offset) in order to trigger water delivery. No water was delivered if mice did not lick the water spout during the response time interval. If mice received a water reward for >80% of trials across 100 trials, they proceeded to the next step.

#### 5. Behavioral shaping step three – full task training

At this step, mice were presented with both match and non-match trials. The timing of the response window was set to 0.5 – 1.5 s from the second odor offset. The delay duration between two odor presentations was set to 3 s at the first session of this step. The delay duration was increased by 0.5 s if mice achieved 70% or more correct performance in the previous behavioral sessions.

Trial type (match/non-match) for each trial was chosen pseudorandomly so that mice did not receive rewards from same trial types in four or more consecutive trials. We also implemented a bias correction strategy. If a choice bias could be predicted from the response history of the past three trials, the trial type of the coming trial was chosen so that the biased response would be incorrect. Otherwise, match and non-match trials were presented with equal probability. For this step, odor mixtures sharing a component (AC and BC) or an odor mixture and one of its component odors (AB and A) were not presented in the same trials to avoid confusing animals. Once mice showed >70% correct performance across 100 trials with 5 sec delay, they were ready to be used to test perceptual similarities between odors of interest.

#### 6. Testing of perceptual similarity

The task was performed as in shaping step three, except that all combinations of first and second odors were used. Delay duration was fixed at 5 s, unless the effect of delay duration was being investigated (**Fig. 3b**, 3 s and 5 s). The first five trials of each session were match trials to encourage mice to lick water spouts, and these trials were excluded from all analyses. The number of mice, sessions, and trials of each data set are summarized in Table 2.

**Table 2.**
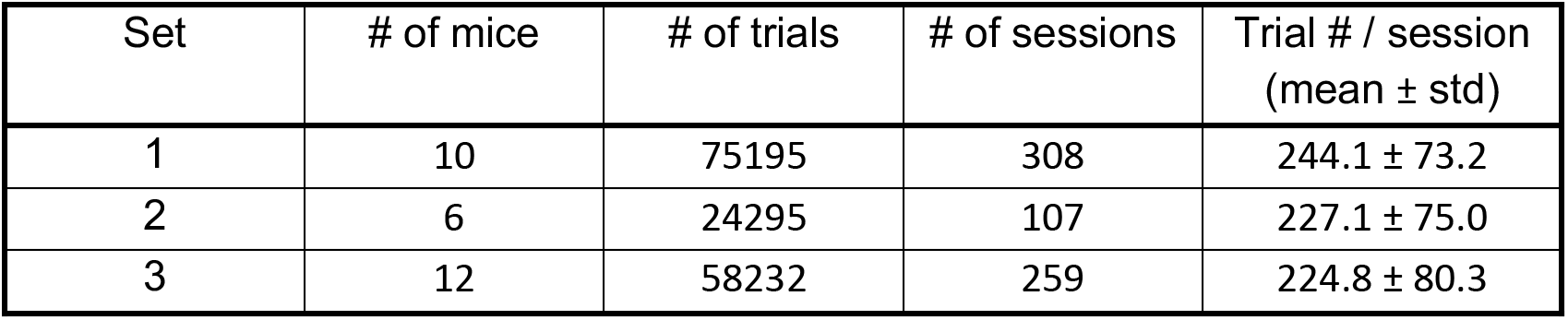
Trial statistics of different odor sets

| Set | # of mice | # of trials | # of sessions | Trial # / session (mean ± std) |
| --- | --- | --- | --- | --- |
| 1 | 10 | 75195 | 308 | 244.1 ± 73.2 |
| 2 | 6 | 24295 | 107 | 227.1 ± 75.0 |
| 3 | 12 | 58232 | 259 | 224.8 ± 80.3 |

### Quantification of perceptual distances

The perceptual distance between a pair of odors was calculated based on the fraction of non-match trials where mice gave the correct response (no-go choice). The probability of a no-go choice for a given odor pair was normalized by the lapse rate for individual odors, i.e. the probabilities of a no-go choice for match trails (see Eq 1). For a reliable estimate of perceptual distances, it is desirable to have at least 100 trials for each odor pair. The transformation from no-go probabilities to the distance measure makes all diagonal elements of the distance matrix take a value of 0. Some non-match odor pairs may take a negative value due to a small number of trials. The negative values were replaced by 0.001 to keep all non-match odor pairs distinct from self-comparison (match trials), which correspond to zero distance.

### Multidimensional scaling

We used non-classical metric multidimensional scaling (MDS) to calculate a low-dimensional representation of odor perceptual space [32, 33]. MDS was performed using the Python scikit-learn package (sklearn.manifold.MDS). MDS takes a pairwise distance matrix as input and returns coordinates of data points in low dimensional space, such that pairwise distances in the input are maximally conserved. Non-classical metric MDS was applied to the matrix representing pairwise perceptual distance between odor pairs. We used a scaling dimension of three for visualization.

### Statistics

Perceptual distances between odor mixtures were compared (**Fig. 2d**) using Wilcoxon signed-rank test. First, the probability of a no-go choice was calculated for each trial type (match, A-B, AC-BC and A-AB) separately for individual mice. Comparisons were performed for pairs (A-B, AC-BC) and (A-B, A-AB) by treating mice as repeated measures. *p*-values were corrected for multiple comparisons with the Bonferroni method.

For the comparison of distances between acids and those between acids and non-acids (**Fig. 2f**), we first calculated perceptual distances across all odor pairs by pooling trials from all animals. Then, we classified the perceptual distances into two groups: distances between acids, and distances between acids and non-acids. Odor pairs including binary mixtures of both acids and non-acids were excluded from this analysis. The difference between these groups was tested with the Mann–Whitney U test.

To quantify the effects of odor identity and other behavioral variables, we fit logistic regression model [36]:

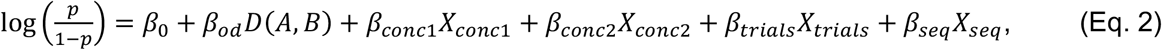

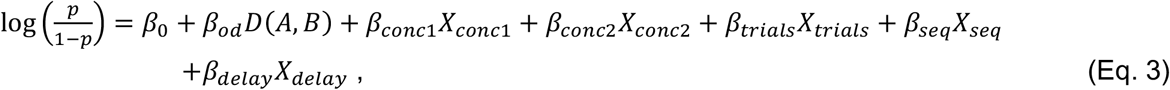

where *p* is probability of no-lick responses; *D(A,B)* is the distance between odors A and B defined in (Eq.1), and calculated separately for each mouse; *X_conc1_* and *X_conc2_* are concentrations of the 1^st^ and 2^nd^ odor in the trail: *X_conc1_,X_conc1_*= 0 *or* 1 for the low or high concentrations; *X_trials_* is a phase of data collection, *X_trials_*= 0 *or* 1 for the 1^st^ vs 2^nd^ half of total set of trials; *X_seq_* is a sequence of odor presentation, *X_seq_*= −1 *or* 1 for A->B and B->A odor presentations in nonmatch trials and *X_seq_*= 0 for match trials. Assignment of values of A->B and B->A sequencies were randomized. *X_delay_* is a delay between sequential odor presentation in one trial, *X_delay_*=0 *or* 1, for 3 s and 5 s delays. Regression coefficients were calculated by averaging results from 50 different random implementations of sequence assignments.

To compare the amplitude of different independent variables, these were standardized so that mean and standard deviation of each variable becomes 0 and 1 respectively. We fit these regression models using statsmodels package in Python [37]. We compared absolute value of regression coefficients using One-way ANOVA and post hoc tests corrected for multiple comparisons (Tukey’s Honestly Significant Difference test).

To illustrate the relationship between the left-hand side of (Eq. 2) or (Eq. 3) and the distance metric, we grouped trials based on the range of values of *D(A,B)*. For each range of *D(A,B)*, we calculated the 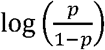 that appears in (Eq. 2) and (Eq. 3).

To calculate the Pearson correlation coefficients of perceptual distance matrices of mouse pairs (**Fig. 3c**), we first we calculated a distance matrix for each mouse as described above. Then, Pearson correlation coefficients between a pair of distance matrices was calculated using only the off-diagonal elements of each matrix. Simulated perceptual distance matrices were created as follows. First, we calculated a matrix of probabilities of no-go choices *(P_all_*) by pooling trials from all mice. The simulated probability of a no-go choice *P_m_(i,j)* for an odor pair (i, j) in mouse-*m* was calculated using a sample from binomial distribution *B(n,p)*.

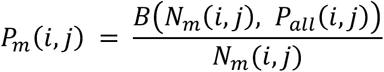

where *N_m_(i,j)* is the number of trials in which mouse m received an odor pair (i, j), *P_all_(i,j)* is the probability of a no-go choice for an odor pair (i, j), calculated from data across all mice. The simulated probability of no-go choice was converted to perceptual distance (Eq. 1) and correlation coefficients between pairs of mice were calculated from simulated perceptual distance matrix for each mouse. The simulated perceptual distance was calculated 1000 times with different random seeds and the average Pearson correlation of these 1000 samples were used as the simulated correlation.

## Acknowledgments

We thank S. Toole, D. Demeterfi, Q. Stewart, and L. Shukla for assistance with building and running experiments. We also thank J. Harvey, J. Mainland, and R. Gerkin for helpful comments. **Funding**: This work was supported by the NIH BRAIN Initiative Grant U19NS112953.

## Author contributions

H.N. and D.R. conceived and designed the experimental, computational and modeling approaches;

H.N. performed experiments and data analyses; H.N. and D.R. wrote the manuscript.

## Data and materials availability

Datasets and analysis code will be available upon a request

---

**Extended figure 1.**
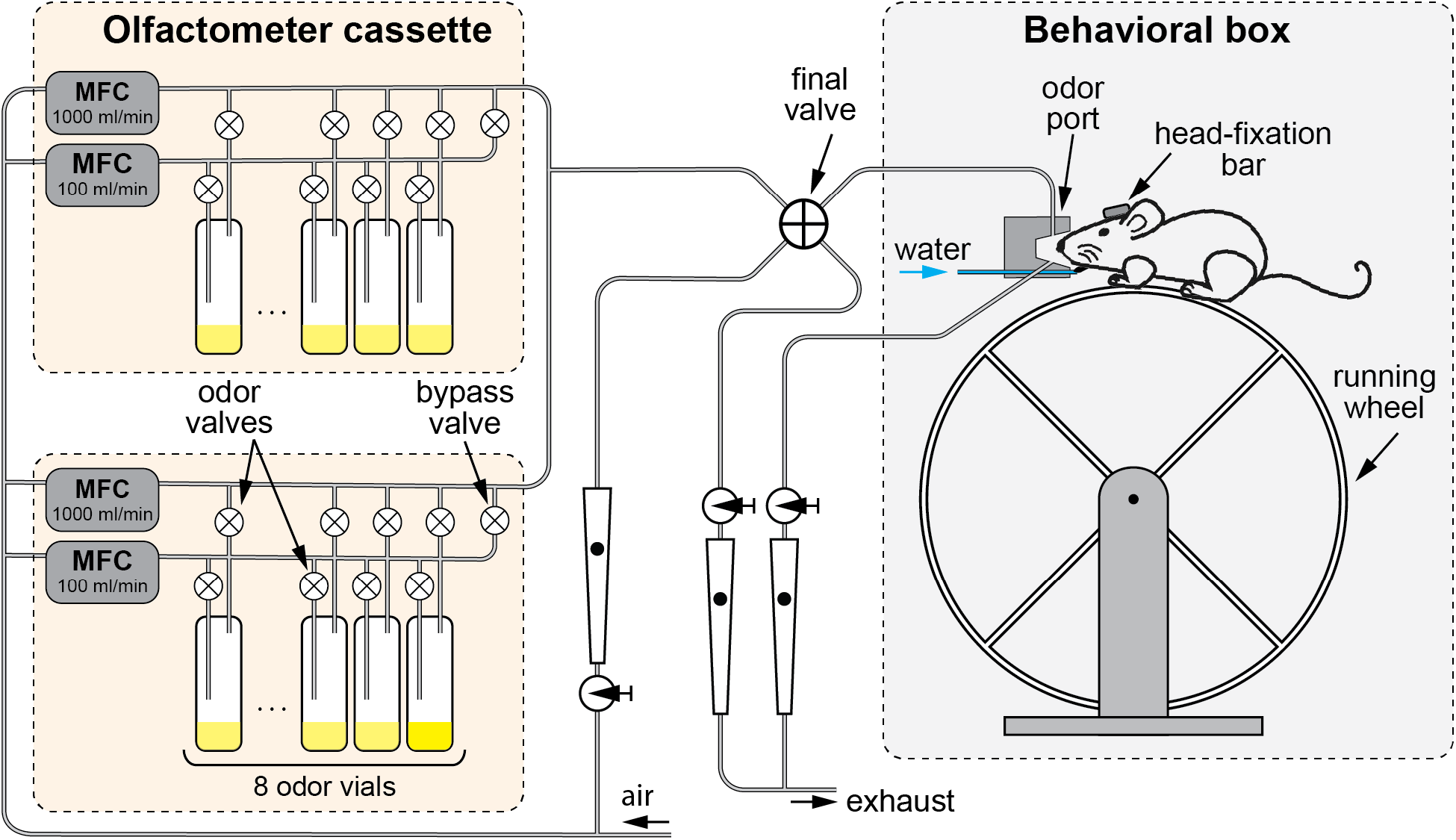
Experimental setup. *Left:* the odor delivery system. Odors are delivered using a two-cassette air dilution olfactometer. Each cassette has 8 odor vials and two Mass Flow Controllers (MFCs) for flow ranges of 0-1000 ml/min and 0-100 ml/min. To prepare an odor a pair of valves for a single odor vial is opened and an odorized air flow (1000 ml/min) is first directed to the exhaust via the final valve. Clean air of the same flow is delivered to the odor port. After approximately 1 sec of the flow stabilization, the final valve redirects the odorized flow to the odor port and the clean air to the exhaust. The concertation is controlled by a ratio of the MFC flows. To deliver a binary mixture, two vials from different cassettes are opened simultaneously. The air is continuously pumped away from the odor port with the same air flow rate. *Right:* behavioral setup. A mouse is positioned on a freely rotating running wheel with its head fixed. The mouse snout is placed in the odor port. The water delivery spout is positioned below the odor port opening.

**Extended figure 2.**
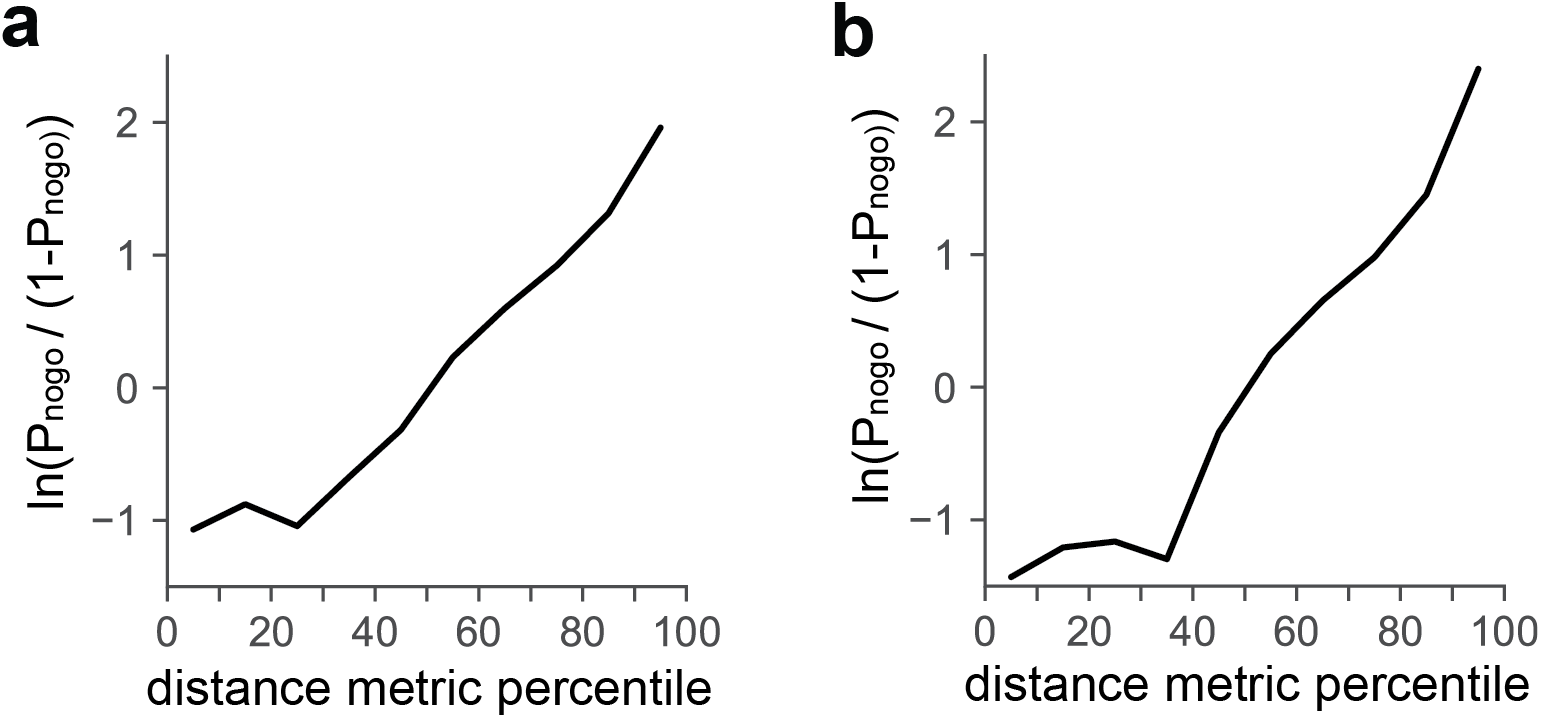
Relationship between distance metric and logit of probability of nogo choice. **a.** plot showing the dependence of logit(*P_nogo_*) (left-hand side of (Eq. 2)) on distance metric between odor pairs. x-axis represents percentile of distance metric across all odor pairs. **b.** same as in a for (Eq.3)

## References

1. Johnson, K.O., S.S. Hsiao, and T. Yoshioka, Neural coding and the basic law of psychophysics. Neuroscientist, 2002. 8(2): p. 111–121.

2. Shepard, R.N., The Analysis of Proximities - Multidimensional-Scaling with an Unknown Distance Function. 2. Psychometrika, 1962. 27(3): p. 219–246.

3. Shepard, R.N., Multidimensional scaling, tree-fitting, and clustering. Science, 1980. 210(4468): p. 390–8.

4. Shepard, R.N., Approximation to uniform gradients of generalization by monotone transformations of scale. Stimulus generalization, 1965: p. 94–110.

5. Ghirlanda, S., et al., A century of generalization. Animal Behaviour, 2003. 66(1): p. 15–36.

6. Guttman, N. and H.I. Kalish, Discriminability and stimulus generalization. Journal of Experimental Psychology, 1956.. 51(1): p. 79–88.

7. Buzsaki, G., et al., Pattern and inhibition-dependent invasion of pyramidal cell dendrites by fast spikes in the hippocampus in vivo. Proc Natl Acad Sci U S A, 1996. 93(18): p. 9921–5.

8. Svoboda, K., et al., In vivo dendritic calcium dynamics in neocortical pyramidal neurons. Nature, 1997. 385(6612): p. 161–5.

9. Chong, E., et al., Manipulating synthetic optogenetic odors reveals the coding logic of olfactory perception. Science, 2020. 368(6497).

10. Emiliani, V., et al., All-Optical Interrogation of Neural Circuits. J Neurosci, 2015. 35(41): p. 13917–26.

11. Gill, J.V., et al., Precise Holographic Manipulation of Olfactory Circuits Reveals Coding Features Determining Perceptual Detection. Neuron, 2020. 108(2): p. 382–393 e5.

12. Packer, A.M., et al., Simultaneous all-optical manipulation and recording of neural circuit activity with cellular resolution in vivo. Nat Methods, 2015. 12(2): p. 140–6.

13. Zhang, F., et al., Multimodal fast optical interrogation of neural circuitry. Nature, 2007. 446(7136): p. 633–9.

14. Mamlouk, A.M. and T. Martinetz, On the dimensions of the olfactory perception space. Neurocomputing: An International Journal, 2004: p. 1019–1025.

15. Meister, M., On the dimensionality of odor space. eLife, 2015. 4: p. e07865.

16. Zhou, Y., B.H. Smith, and T.O. Sharpee, Hyperbolic geometry of the olfactory space. Sci Adv, 2018. 4(8): p. eaaq1458.

17. Dravnieks, A., A.C.E.-o.S.E.o. Materials, and P.S.E.-o.O. Profiling, Atlas of odor character profiles. 1985: ASTM.

18. Koulakov, A.A., et al., In search of the structure of human olfactory space. Frontiers in systems neuroscience, 2011. 5: p. 65.

19. Wise, P.M., M.J. Olsson, and W.S. Cain, Quantification of odor quality. Chem Senses, 2000. 25(4): p. 429–43.

20. Keller, A., et al., Predicting human olfactory perception from chemical features of odor molecules. Science, 2017. 355(6327): p. 820–826.

21. Secundo, L., et al., Individual olfactory perception reveals meaningful nonolfactory genetic information. Proc Natl Acad Sci U S A, 2015. 112(28): p. 8750–5.

22. Snitz, K., et al., Predicting odor perceptual similarity from odor structure. PLoS Comput Biol, 2013. 9(9): p. e1003184.

23. Cleland, T.A., et al., Behavioral models of odor similarity. Behav Neurosci, 2002. 116(2): p. 222–31.

24. Etkin, M. and M.R. D’Amato, Delayed matching-to-sample and short-term memory in the capuchin monkey. Journal of Comparative and Physiological Psychology, 1969. 69(3): p. 544–549.

25. Grant, D.S., Proactive interference in pigeon short-term memory. Journal of Experimental Psychology: Animal Behavior Processes, 1975.. 1(3): p. pp.

26. Koopman, S.E., B.Z. Mahon, and J.F. Cantlon, Evolutionary Constraints on Human Object Perception. Cogn Sci, 2017. 41(8): p. 2126–2148.

27. Otto, T. and H. Eichenbaum, Complementary roles of the orbital prefrontal cortex and the perirhinal-entorhinal cortices in an odor-guided delayed-nonmatching-to-sample task. Behav Neurosci, 1992. 106(5): p. 762–75.

28. Liu, D., et al., Medial prefrontal activity during delay period contributes to learning of a working memory task. Science (New York, NY), 2014. 346(6208): p. 458–463.

29. Taxidis, J., et al., Differential Emergence and Stability of Sensory and Temporal Representations in Context-Specific Hippocampal Sequences. Neuron, 2020. 108(5): p. 984–998 e9.

30. Wu, Z., et al., Context-Dependent Decision Making in a Premotor Circuit. Neuron, 2020. 106(2): p. 316–328 e6.

31. Shepard, R.N., Toward a universal law of generalization for psychological science. Science, 1987. 237(4820): p. 1317.

32. Borg, I. and P.J.F. Groenen, Modern Multidimensional Scaling Theory and Applications. 1997.

33. Kruskal, J.B., Nonmetric multidimensional scaling: A numerical method. Psychometrika, 1964.. 29(2): p. pp.

34. Kruskal, J.B., Multidimensional scaling by optimizing goodness of fit to a nonmetric hypothesis. Psychometrika, 1964.. 29(1): p. pp.

35. Ferreira, V., Revisiting psychophysical work on the quantitative and qualitative odour properties of simple odour mixtures: a flavour chemistry view. Part 2: qualitative aspects. A review. Flavour and Fragrance Journal, 2012. 27(3): p. 201–215.

36. James, G., Witten, D., Hastie, T., & Tibshirani, R., Classification, in An Introduction to Statistical Learning. 2013, Springer: New York. p. 127–174.

37. Seabold, S. and J. Perktold, Statsmodels: Econometric and statistical modeling with python. Proceedings of the 9th Python in Science Conference, 2010.

